# The interaction of dispersal, species sorting, and priority effects, explains a unimodal response of β-diversity to immigration rates

**DOI:** 10.64898/2025.12.31.697192

**Authors:** Esteban Ortiz, Rodrigo Ramos-Jiliberto, Matías Arim

**Affiliations:** Departamento de Ecología y Gestión Ambiental. Centro Universitario Regional Este (CURE), Universidad de la República, Uruguay; GEMA Center for Genomics, Ecology & Environment. Universidad Mayor, Chile

## Abstract

A dominant perspective in metacommunity theory is that dispersal consistently homogenizes species composition across local communities. However, recent studies have reported a direct relationship between dispersal rate and β-diversity, which has been attributed to the influence of priority effects and species sorting. The underlying mechanisms by which dispersal can either increase or decrease β-diversity remain unclear, hindering our understanding of metacommunity assembly and the processes driving biodiversity patterns across time and space. We theoretically evaluated how purely random priority effects, priority effects conditioned by species dispersal abilities, and species sorting shape the β-diversity–dispersal relationship in metacommunities open to exogenous dispersal (i.e., immigration) from a regional species pool. Our results show that the three evaluated processes foster community differentiation at intermediate exogenous dispersal rates when dispersal between communities (i.e. endogenous dispersal) is taking place. Additionally, the interaction between endogenous and exogenous dispersal homogenizes communities at low and high exogenous dispersal rates, determining a unimodal response of β-diversity to increasing exogenous dispersal rates. Our findings contribute to a deeper mechanistic understanding of metacommunity assembly and offer a coherent explanation for previously observed empirical discrepancies, thereby challenging the traditional view of dispersal as a purely homogenizing process.

## Introduction

Detecting the main mechanisms that determine biodiversity patterns remains a major goal in ecology (Leibold and Chase 2018). Metacommunity theory has provided a robust framework that combines the interaction between local and regional processes to explain the emergence of diversity across multiple spatial scales (Leibold et al. 2004; Thompson et al. 2020; Suzuki and Economo 2021). The degree of dissimilarity among a set of communities is captured by β-diversity, which is conceived as an important determinant of biomass production and ecosystem functioning in heterogeneous environments (van der Plas et al. 2023). In this context, the β-diversity–dispersal relationship has been a major focus of attention. Dispersal has traditionally been conceptualized as a homogenizing process that increases similarity in species composition among communities (Loreau and Mouquet 1999; Loreau 2000; Soininen et al. 2007; Mouquet and Loreau 2003; Grainger and Gilbert 2016; Heino and Tolonen 2017; Gu et al. 2023). Foundational theoretical studies predicted that β-diversity decreases monotonically with dispersal intensity among communities (Mouquet and Loreau 2003; Leibold et al. 2004; Soininen et al. 2007), and a large body of field and experimental studies has remarkably supported this prediction (Grainger and Gilbert 2016; Catano et al. 2017; Simon-Lledó et al. 2025). However, recent empirical (Werner et al. 2007; Catano et al. 2017; Ojima and Jiang 2017; Vannette and Fukami 2017; Borthagaray et al. 2025) and theoretical (Lu 2021; Borthagaray et al. 2025) studies have evidenced more complex relationships between dispersal and β-diversity, including cases of positive associations.

The detection of complex (i.e., positive or non-monotonic) relationships between β-diversity and dispersal has emerged under conditions in which: (i) dispersal rates and local conditions are naturally allowed to vary during metacommunity assembly; (ii) interspecific differences in dispersal abilities and local performance are explicitly considered; and/or (iii) metacommunities experience immigration from an external source of species (i.e., an external species pool). A more detailed treatment of these mechanisms is provided in the Discussion. Within this framework, the positive effects of dispersal on β-diversity have been ultimately attributed to the operation of priority effects (Chasse 2003; Fukami 2015; Vannette and Fukami 2017) and species sorting (Cottenie and De Meester 2004; Werner et al. 2007; Gianuca et al. 2017), both of which are widely recognized as important determinants of community dissimilarity. Low to intermediate dispersal rates can enhance the operation of priority effects and species sorting by increasing the diversity of species that act as potential colonizers of communities (Chase 2003; Fukami 2015; Vannette and Fukami 2017) and by allowing a broader set of species to track regional environmental heterogeneity (Heino 2011; Soininen 2014; Alahuhta et al. 2014). These processes facilitate the emergence of alternative species compositions among local communities, thereby increasing β-diversity. In addition, immigration from a regional species pool supplies metacommunities with a continuous influx of individuals exhibiting diverse traits, which serve as the source of diversity upon which priority effects and species sorting can act (Cottenie and De Meester 2004; Leibold et al. 2004; Gianuca et al. 2017). Despite this, the role of dispersal in promoting community differentiation has been systematically underestimated in empirical studies (Logue et al. 2011; Grainger and Gilbert 2016; Vannette and Fukami 2017) by ignoring these sources of variation. Consequently, we still lack a comprehensive understanding of how dispersal explicitly modulates the ultimate determinants of community dissimilarity, namely species sorting and stochastic dynamics driven by priority effects.

In metacommunities open to a regional species pool (Figure 1), we therefore expect a humped relationship between β-diversity and the rate of immigration into the metacommunity (hereafter referred to as *exogenous or external dispersal*, following Fukami 2005) under three alternative scenarios: purely random priority effects (H_1_), priority effects conditioned by species dispersal ability (H_2_), and species sorting (H_3_). Low exogenous dispersal limits the diversity of species reaching each local patch, thereby constraining community differentiation. Under this scenario, communities tend to be dominated by a reduced set of pioneer species through stochastic priority effects (H_1a_), by priority effects associated with highly dispersive species (H_2a_), or by a reduced set of locally adapted species via species sorting (H_3a_). In all cases, dispersal among local communities (hereafter referred to as *endogenous or internal dispersal,* following Fukami 2005) enables this limited set of immigrant species to spread within the metacommunity, further diminishing β-diversity. In contrast, at intermediate exogenous dispersal rates, the diversity of species arriving from the regional pool increases, strengthening both priority effects and species sorting. Community differentiation may then arise through alternative dominance by different pioneer species (H_1b_), which may also exhibit contrasting dispersal abilities (H_2b_), or through species sorting resulting from the differential selection of species with distinct traits (H_3b_). Finally, at high exogenous dispersal rates, mass effects homogenize local communities, overriding both priority effects (H_1c_, H_2c_) and species sorting (H_3c_), and thereby reducing β-diversity.

**Figure 1.**
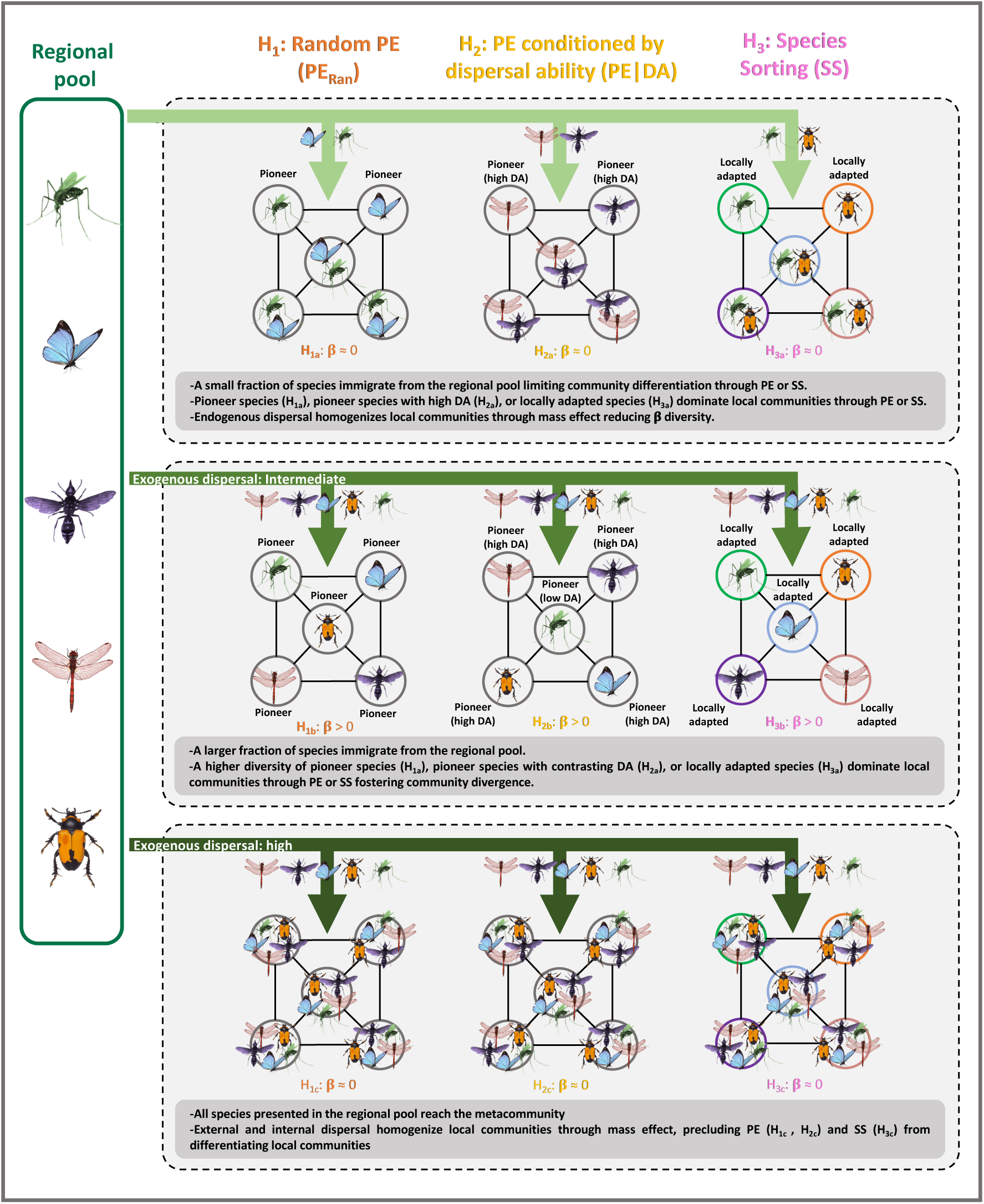
Hypotheses predicting a humped relationship between β-diversity and dispersal from the regional species pool (i.e. exogenous dispersal), driven by three mechanisms: random priority effects (H_1_), priority effects conditioned by species dispersal ability (H_2_), and species sorting (H_3_).

In this study, we theoretically test these hypotheses by analyzing the effects of species arrival time, interspecific variation in dispersal abilities, and local performance on the relationship between β-diversity and dispersal. We employed a lottery model to simulate the dynamics of metacommunities connected to a regional species pool and to quantify β-diversity patterns along an exogenous dispersal gradient, under scenarios of purely random priority effects (H_1_), priority effects conditioned by differences in dispersal ability (H_2_), and species sorting (H_3_) (Figure 1).

## Methods

### The lottery-based metacommunity model

Metacommunities were assembled using a lottery model (Hubbell 2001; Borthagaray et al. 2014; Worm and Tittenson 2018; Cunillera-Montcusí et al. 2021; Ortiz et al. 2023a; Borthagaray et al. 2025), which simulates the dynamics of a set of local communities that are interconnected and linked to a regional species pool through endogenous and exogenous dispersal, respectively (Figure 2). In addition, our model incorporates interspecific variation in dispersal ability and environmental filtering during the assembly process of the metacommunities. In its simplest form, the model operates in two stages (Figure 2). The first stage involves community filling through coalescent dynamics. Communities are initially colonized by a different individual randomly sampled from the regional species pool (Figure 2a, Step 1). They are then filled with J individuals each, progressively adding new individuals randomly selected either from the regional species pool (with probability *m.pool*), from neighboring communities (with probability *m.meta*), or from the focal community through local recruitment (with probability 1 – *m.pool* – *m.meta*) (Figure 2a, Step 2) (Worm and Tittenson 2018; Cunillera-Montcusí et al. 2021; Borthagaray et al. 2023a, b; Ortiz et al. 2023a; Borthagaray et al. 2025). Once all communities have reached J individuals each, lottery dynamics start (Figure 2b) (Hubbell 2001; Worm and Tittenson 2018; Cunillera-Montcusí et al. 2021; Ortiz et al. 2023a). At each time step in each local community, a single individual is randomly removed (Figure 2b, Step 1) to simulate a local death and is replaced by a recruit randomly selected either from the regional pool, from neighboring communities or from the local community itself, with the same probabilities as described above (Figure 2b, Step 2). This procedure is repeated for N_it_ iterations until the dynamics stabilize. The model simulates the dynamics of all local communities simultaneously, such that colonization, dispersal, mortality, and recruitment occur concurrently across the metacommunity.

**Figure 2.**
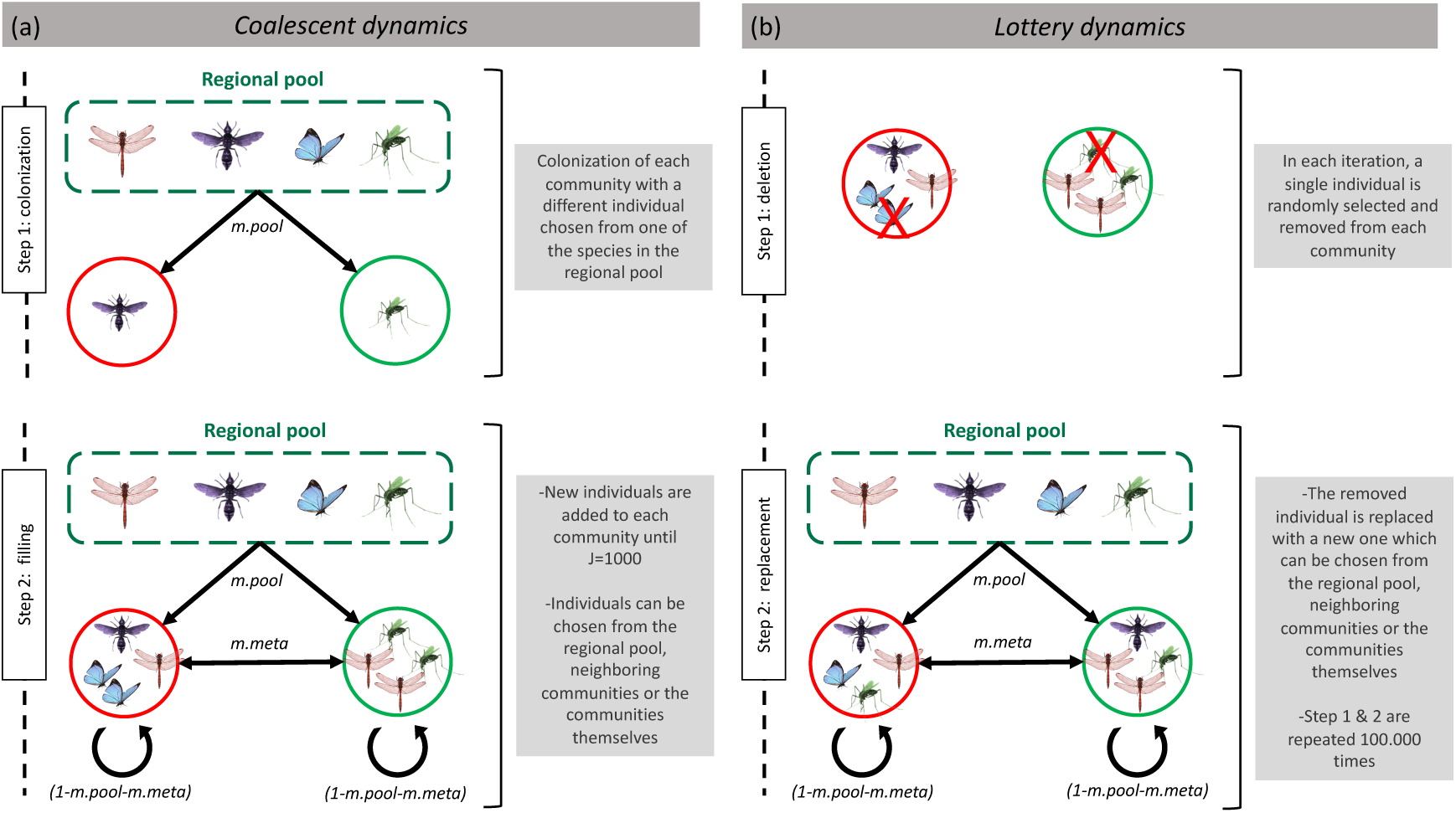
Lottery-based metacommunity model pseudocode. Metacommunities are assembled by recruiting individuals from the regional species pool, from neighboring communities, and from the focal local communities (local recruitment).

### Evaluation of mechanisms

#### Hypothesis 1: Random priority effects

To evaluate how purely random priority effects (H_1_) modulate the relationship between β-diversity and dispersal, we implemented a neutral version of the model in which species are equivalent in both dispersal ability and local performance. In this scenario, the probability of recruiting an individual of species *i* from the regional pool (*P*_*i*,*pool*_), or from a neighboring community *q* (*P*_*i*,*q*_) in each time interval (i.e., iteration) is determined exclusively by the relative abundance of species *i* at the source, *f*_*i*,*pool*_ or *f*_*i*,*q*_, and by the probability that an individual disperses from the source to the focal patch. These dispersal probabilities are given by *m.pool* and *m.meta,* such that

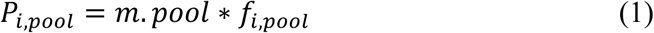

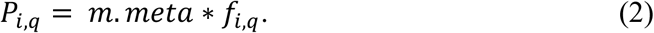

The probability of local recruitment of an individual of species *i* from any focal community *p* (*P*_*i*,*p*_), and the probability of local death (*D*_*i*,*p*_) are defined as:

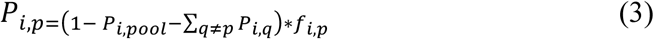

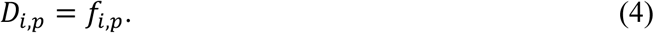

#### Hypothesis 2: Priority effects conditioned by dispersal ability

To evaluate the effect of interspecific differences in dispersal ability on the β-diversity–dispersal relationship, each species *i* was assigned a dispersal ability value *DA*_*i*_, as: *DA*_*i*_ = *x*_*i*_^10∗*Q*^. Here, *x*_*i*_ is a value uniformly drawn from the interval [0,1], and *Q* is a parameter that determines the magnitude of the interspecific differences in *DA*_*i*_(Supplementary Material, Figure S1a and S2a). Dispersal ability values were subsequently standardized and used to compute species-specific probabilities of dispersal and local recruitment such that larger *DA*_*i*_ values correspond to larger dispersal abilities. Now, in each iteration, the probability of recruiting an individual of species *i*, either from the regional pool (*P*_*i*,*pool*|*DAi*_) or from any adjacent community *q* (P_*i*,*q*|*DAi*_), is defined by its dispersal ability value as:

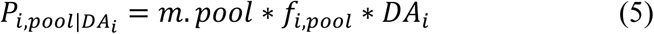

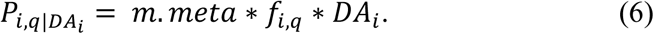

Consequently, the probability of recruiting an individual of species *i* from any focal community *p* (*P*_*i*,*p*_) is now:

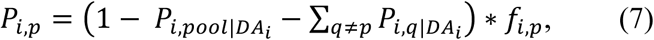

and the probability of local death (*D*_*i*,*p*_) remains the same as in eqn. (4).

#### Hypothesis 3: Species sorting

To incorporate the effect of species sorting into the dynamics, a distinct environmental and trait value distribution were assigned to each local community and species respectively (Supplementary Material, Figure S1b). The mean value *μ* of each distribution was placed within the interval [0,1] with standard deviation *σ*. The degree of overlap between the trait value distribution of species *i* and the filter value distribution of community *p* (i.e., the filter–trait match —*MFT*_*i*,*p*_—) determines the local performance of species *i* in community *p*. That is, the larger the *MFT*_*i*,*p*_, the stronger the positive local selection acting on species *i*. As in the previous scenario, *MFT*_*i*,*p*_ values are also standardized to calculate the probabilities of local recruitment and death. Under this scenario, the probabilities of recruiting a new individual of species *i* per time step, either from the regional pool (*P*_*i*,*pool*_) or from an adjacent community *q* (*P*_*i*,*q*_), obey eqns (1) and (2). However, the probabilities of local recruitment (*P*_*i*,*p*|*MFT*_*i*,*p*_) and death (*D*_*i*,*p*|*MFT*_*i*,*p*__) are now defined by the *MFT*_*i*,*p*__ value, as:

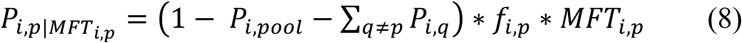

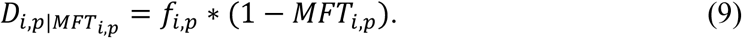

### Data analysis

We simulated metacommunities composed of five interconnected local communities, each with *J* = 1000 individuals, connected to a regional species pool containing 100 species. Simulations were run for varying exogenous dispersal rates, *m.pool* in the range [0, 0.5], for different values of endogenous dispersal among communities, *m.meta*, with 10 replicates per parameter combination. In addition, we ran simulations considering different magnitudes of interspecific differences in dispersal ability and local environmental filters (Supplementary Material, Figure S2). Lottery dynamics were run for 100,000 time steps in the scenarios of random priority effects (H_1_) and priority effects conditioned by dispersal ability (H_2_), and 50,000 time steps in the species sorting scenario (H_3_). These simulation lengths were sufficient to allow diversity to stabilize and to eliminate transient dynamics (Supplementary Material, Figure S3).

At the end of each simulation, we calculated β-diversity as the mean unweighted Jaccard dissimilarity among all pairs of local communities. We also calculated α-diversity as the mean species richness across communities, and γ-diversity as the total species richness of the metacommunity. To assess how random priority effects (H_1_), priority effects conditioned by species dispersal ability (H_2_), and species sorting (H_3_) shape community assembly and species abundance, we quantitatively evaluated how species arrival times to local communities (*AT*_*i*,*p*_), as well as their dispersal abilities (*DA*_*i*_) and local performance (*MFT*_*i*,*p*_) determined their final abundances along the *m.pool* gradient. All analyses were performed using R software v.4.0.4 (R Core Team 2021).

## Results

### β-diversity–dispersal relationship

β-diversity exhibited complex responses along the exogenous dispersal gradient (*m.pool*), with patterns strongly modulated by the endogenous dispersal rate (*m.meta*) (Figure 3). When *m.meta* was null, β-diversity decreased monotonically with increasing *m.pool* (Figure 3a). In contrast, β-diversity displayed a unimodal relationship with *m.pool* at low (Figure 3b), intermediate (Figure 3c), and high (Figure 3d) levels of *m.meta*. This qualitative pattern was consistent across the three simulated scenarios, although it was more pronounced when interspecific differences in dispersal ability and local selection were incorporated. Our results were robust to the choice of the β-diversity metrics (see supplementary material, Figure S4). Additionally, α- (Figure 3e– h) and γ-diversity (Figure 3i–l) consistently increased with the magnitude of exogenous dispersal. See Supplementary Material Figure S5 for the results of α-, β-, and γ-diversity across the parameter space.

**Figure 3.**
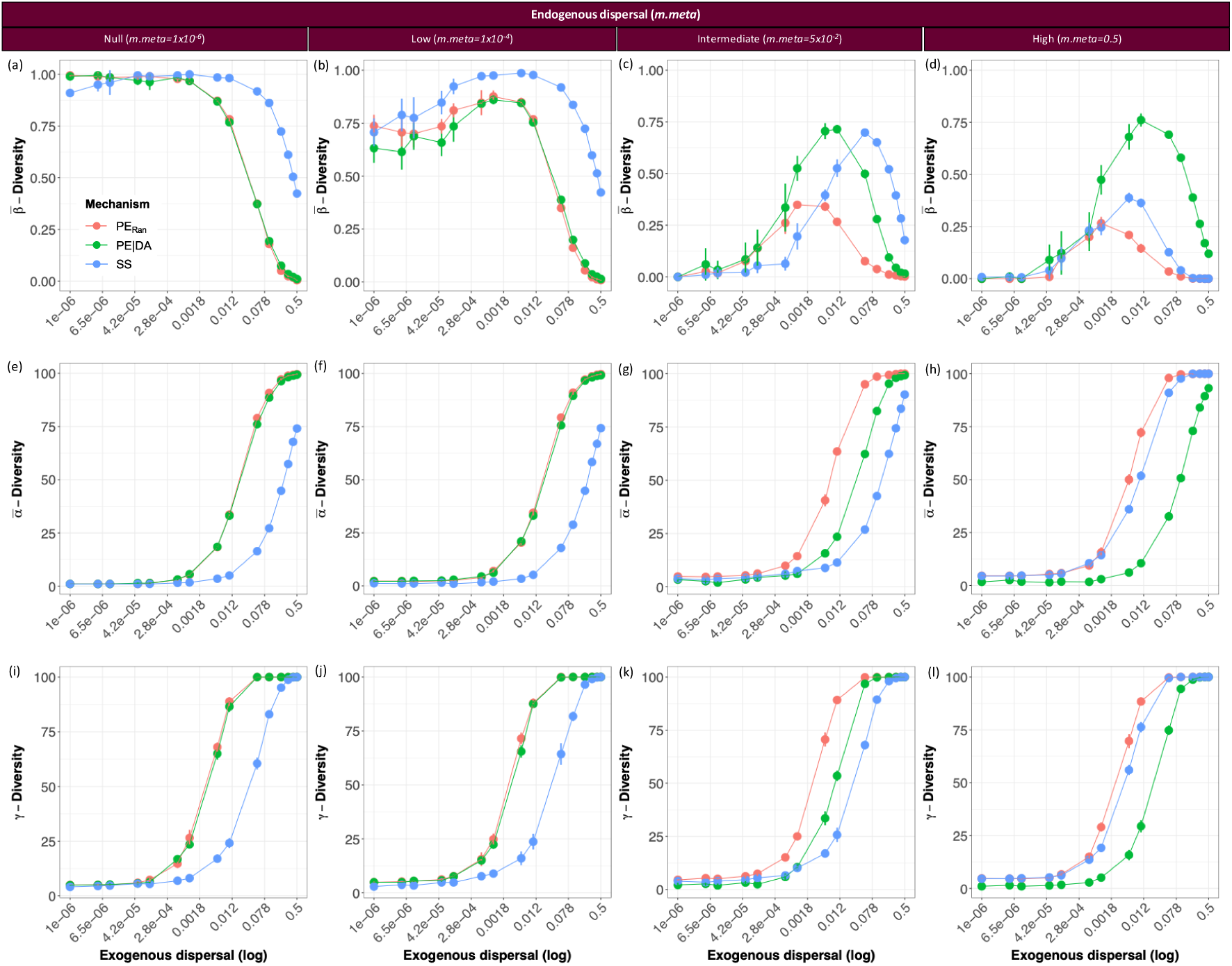
Responses of mean β-diversity (a-d), mean α-diversity (e-h), and γ-diversity (i-l) to the exogenous dispersal rate (*m.pool*) under varying levels of endogenous dispersal rate (*m.meta*). Symbols represent the mean and 95% confidence interval, calculated from 10 replicates.

### Evaluation of mechanisms

Our previous results (Figure 3) show that high exogenous dispersal rates reduce β-diversity across all scenarios, whereas endogenous dispersal rates reduce β-diversity when exogenous dispersal is low. However, β-diversity remains high despite elevated endogenous dispersal when exogenous dispersal is not too strong. Below, we analyze the role of the three evaluated hypotheses in explaining the rise and decline in β-diversity with increasing exogenous dispersal rates. Specifically, we assess the efficacy of the three proposed alternative mechanisms by which β-diversity resists the homogenizing effect of endogenous dispersal at moderate exogenous dispersal levels.

#### Hypothesis 1: Random priority effect

The arrival times of species to local communities (*AT*_*i*,*p*_) determined species abundances (*N*_*i*,*p*_) and community dissimilarity via priority effects (Figure 4a). At low and intermediate exogenous dispersal rates, early-arriving attained markedly higher abundances than late-arriving species. In contrast, high external dispersal rates equalized abundance levels between early- and late-arriving species, canceling the impact of priority effects on species abundances.

**Figure 4.**
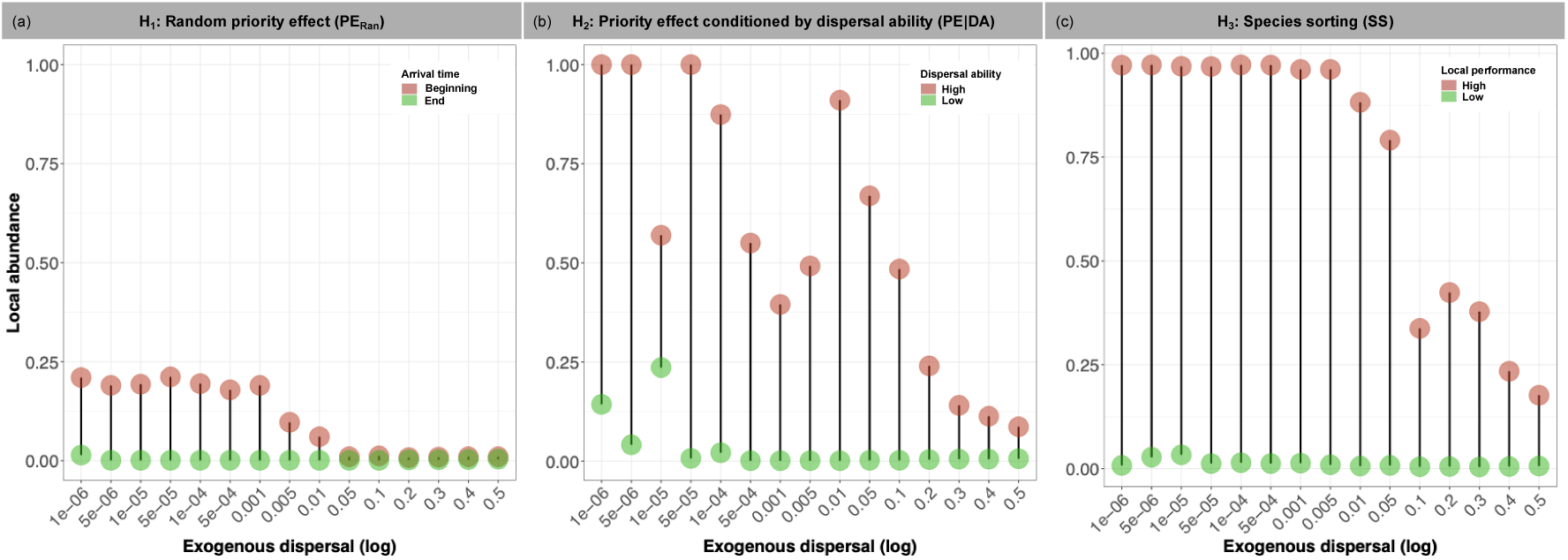
Effect of species arrival time (*AT*sub_*i*,*p*_) (a), species dispersal ability (*DA*_*i*_) (b), and species local performance (*MFT*_*i*,*p*_) (c) on local asymptotic species abundance across a gradient of exogenous dispersal rates (*m.pool*), at high endogenous dispersal rates (*m.meta* = 5×10^−2^). Red and green symbols represent species in the earliest and latest 1% of arrivals (a), species in the highest and lowest 1% of dispersal abilities (b), and species in the highest and lowest 1% of local performances.

#### Hypothesis 2: Priority effects conditioned by species dispersal ability

Species dispersal ability (*DA*_*i*_) determined species abundances and community dissimilarity by favoring the early arrival of species with high dispersal rates to the metacommunity (Figure 4b). Species with large *DA*_*i*_ values consistently exhibited higher local abundances than those with low dispersal abilities. Differences in abundance between good and bad dispersers were strongly reduced at high *m.pool* values (see also Supplementary Material, Figure S6).

#### Hypothesis 3: Species sorting

Species local performance (*MFT*_*i*,*p*_) determined species abundances (*N*_*i*,*p*_) and community dissimilarity through species sorting (Figure 4c). Species with large *MFT*_*i*,*p*_values consistently showed higher local abundances than species with low local performance, and these differences were reduced at high *m.pool* values. Furthermore, species abundance was not associated with their arrival times, indicating that species sorting outweighed priority effects during the assembly process (Supplementary Material, Figure S7).

## Discussion

A large body of research has shown that stochasticity in species arrival times to local patches (Lepori and Malmqvist 2009; Fukami 2005, 2015; Vannette and Fukami 2017), interspecific differences in dispersal ability (De Bie et al. 2012; Jones et al. 2015; Hill et al. 2017; Vannette and Fukami 2017; Cunillera et al. 2020; Ortiz et al. 2023a) and local selection driven by heterogeneity in environmental filters (Cottenie and De Meester 2004; Mouquet et al. 2006; Gianuca et al. 2017; Cadotte and Tucker 2017) interact with dispersal, shaping β-diversity. Nevertheless, the potential role of dispersal in promoting community differentiation has been overlooked by most empirical studies by ignoring these sources of variation (Logue et al. 2011; Grainger and Gilbert 2016; Vannette and Fukami 2017). Typically, experimental setups predefine the initial composition of species avoiding communities from being stochastically colonized, consider patches with homogeneous environmental conditions limiting community assembly by species sorting, and eliminate any natural differences in dispersal ability between species (Logue et al. 2011; Grainger and Gilbert 2016; Vannette and Fukami 2017). Additionally, models and experiments often simulate only endogenous dispersal among communities and do not consider metacommunities connected to a regional species pool (Grainger and Gilbert 2016; Logue et al. 2011). Here, we show that these sources of variation can determine a humped relationship between β-diversity and the immigration gradient from the regional pool, but only when sufficient dispersal among communities is considered (Figure 3b–d). This result must be highlighted since the humped trend in β-diversity was observed even under the action of mechanisms representing contrasting biological scenarios (i.e., selective vs. stochastic assembly). Specifically, β-diversity was highest when exogenous dispersal: (i) optimized the number of species that colonized local communities, determining different assembly histories through priority effects (Figure 4a); (ii) allowed the arrival and establishment of species with contrasting dispersal abilities in local communities which would be otherwise dominated only by good dispersers (Figure 4b and Figure S6); and (iii) allowed the arrival of species with different traits to those local communities where they were best adapted, determining community divergence through species sorting (Figure 4c). Furthermore, community homogenization arose from the interaction between exogenous and endogenous dispersal. When exogenous dispersal was low, endogenous dispersal facilitated the spread within the metacommunity of the reduced set of species that arrived from the regional pool. In contrast, high exogenous dispersal homogenized local communities through mass effect surpassing priority effects and species sorting (Fukami 2005, 2015; Gianuca et al. 2017; Loke and Chisholm 2023). Overall, these results are fully consistent with our hypotheses (Figure 1).

Under priority effects, community structure and differentiation result from variations in the arrival times of species to local communities (Fukami 2015). Species arriving at early stages of the assembly process can dominate local communities by monopolizing limiting resources (i.e., niche preemption), reducing the abundance and viability of those species that arrive later during community assembly (Fukami 2015). The importance of colonization history in structuring communities is expected to be particularly pronounced when species present high niche overlap (Vannette and Fukami 2014), and at low to intermediate immigration rates from the regional pool (Chase 2003; Fukami 2005). This provides pioneer species with more time to establish and grow before the arrival of other species, reducing subsequent competitive exclusion driven by differences in local performance. This was the case for our simulations when metacommunities were neutrally assembled (H_1_) (Figure 4a), detecting high β-diversity at intermediate exogenous dispersal. However, priority effects are usually conceived under a source-sink conceptualization, in which species arrive at a set of local, unconnected communities from an external source of immigrants (Fukami 2015). Under this scenario, β-diversity is expected to decrease as immigration rates from the regional pool increase due to species accumulation through mass effect (Chase 2003; Fukami 2015). This prediction is in line with our results when endogenous dispersal was nearly null (Figure 3a). However, by considering the interaction between exogenous and endogenous dispersal, we were able to also detect a positive relationship between β-diversity and dispersal (Figure 3b–d). Future empirical studies are needed aiming at disentangling how biodiversity patterns are modulated by priority effects and the interplay between exogenous and endogenous dispersal in natural and experimental systems.

Priority effects are typically expected when species exhibit similar dispersal abilities (Fukami 2015). If species differ in their dispersal rates, a deterministic sequence in the arrival of species to local communities is expected, being the good dispersers the first ones to arrive, limiting community divergence driven by historical contingency (Fukami 2015). However, our results show that β-diversity can also be increased under scenarios in which species differ in their dispersal abilities (H_2_). At low exogenous dispersal, communities were colonized and dominated by a reduced set of species with high dispersal ability (Figure 4b and Figure S6), which spread to other communities through endogenous dispersal. In contrast, intermediate exogenous dispersal allowed the arrival of a larger set of species with suboptimal dispersal abilities to the metacommunity. The increase in community differentiation can be explained either by historical contingency if the species with suboptimal dispersal abilities are able to colonize and dominate different communities, or by rescue effect (Hanski 1982; Gotelli 1991) since the local populations of these sub-optimal dispersers are expected to receive more recruits. Further investigation is needed in this area, since the interaction between these two mechanisms, and their effect on community diversity may have been overlooked in experimental studies that eliminate the variability in dispersal ability among species (Grainger and Gilbert 2016; Logue et al. 2011).

In agreement with several empirical studies (Cottenie and De Meester 2004; Gianuca et al. 2017), the increment from low to intermediate exogenous dispersal enhanced the role of species sorting in the structuring of communities, whereas at high immigration rates this effect was overridden by mass effects. Communities subject to low and intermediate immigration rates from the regional pool were strongly structured by species sorting, even outweighing priority effects (Figure 4c and Figure S7). This result reinforces the idea that both exogenous and endogenous dispersal play a major role in allowing species to arrive at suitable communities where they are positively selected (Cottenie and De Meester 2004; Leibold et al. 2004; Gianuca et al. 2017), thereby promoting β-diversity. It is important to note that although our model incorporates species sorting driven by environmental filtering (H_3_), it can also be conceptually extended to encompass biotic interaction that could generate differential selection between species across the metacommunity (e.g., interference competition, predation, mutualisms) (García-Girón et al. 2020; Ortiz et al. 2023b; Argueta-Guzmán et al. 2025; Souza et al. 2025). However, an explicit evaluation of the effects of biotic interactions on the β-diversity–dispersal relationship in multitrophic metacommunities is still needed.

The influence of a stable, independent regional pool of species that supplies metacommunities with an external set of species has been highlighted as an important determinant of community assembly (Fukami 2005, 2010; Harrison and Cornell 2008; Borthagaray et al. 2025). However, a concrete interpretation of what the regional pool represents in empirical contexts is not always straightforward. Under our conceptualization, the regional pool of species can represent either a source of immigrants at a biogeographical scale that arrive at a focal metacommunity, or a spatially delimited portion of a metacommunity (e.g., a mainland) that acts as a source of individuals to a subset of local communities within the same metacommunity. In both cases, it is essential that the dynamics within the regional pool are decoupled from those operating within the metacommunity to explore how β-diversity is affected by dispersal (Fukami 2015). Empirical studies have frequently overlooked the importance of incorporating an external source of immigrants by working with closed metacommunities (Grainger and Gilbert 2016; Logue et al. 2011). We draw attention to this issue and highlight that more empirical research is needed to determine how the regional species pool interacts with metacommunity dispersal in shaping community structure and biodiversity patterns.

Our results are consistent with, and complement, a range of empirical and theoretical studies that also challenged the traditional view of dispersal as a purely homogenizing process. In a metacommunity of nectar microbes, dispersal among flowers promoted β-diversity due to strong priority effects when pollinators with different dispersal abilities were considered (Vannette and Fukami 2017). Similarly, pond connectivity—a proxy for dispersal rate—combined with environmental heterogeneity was found to increase β-diversity in amphibian assemblages (Werner et al. 2007). In addition, a lack of evidence for the expected homogenizing effect of dispersal on β-diversity was reported in experiments with protist (Ojima and Jiang 2017) and herbaceous plant communities (Catano et al. 2017) subject to disturbance. By applying occupancy-based models on metacommunities connected to a regional species pool, a humped pattern in β-diversity was associated with endogenous dispersal (but not exogenous dispersal as was our case) (Lu 2021). This pattern emerged when species were constrained in the number of patches they can occupy. The resulting reduction in patch occupancy was attributed to different selective regimes determined by biotic interactions and disturbance intensity, which suppress the occurrence probability of dominant species. It is worth noting that this explanation also applies when exogenous dispersal is considered, as shown in our study. Finally, a U-shaped pattern in fish β-diversity along a connectivity (i.e., dispersal) gradient in riverine metacommunities was both theoretically predicted and empirically observed (Borthagaray et al. 2025). This novel pattern likely emerges from distinct community assembly processes between headwaters and downstream sites. Communities in headwaters are primarily differentiated by ecological drift, whereas those near the outlet are shaped by the influx of species from it. The outlet thus acts as a reservoir of species that enhances β-diversity in nearby communities (exposed to higher exogenous dispersal rates) compared with more distant sites.

Our results can be heuristically interpreted through an analogy with economic systems in which external capital inflows and internal circulation play distinct roles. Exogenous dispersal, analogous to external investment or immigration, primarily increases total system richness, as reflected by monotonic increases in α- and γ-diversity. In contrast, β-diversity depends on how this diversity is redistributed internally. High external inflow homogenizes communities through mass effects, much as strong market integration reduces regional differentiation. Conversely, when external inflow is low, homogenization arises only if internal redistribution is sufficiently strong to spread a limited pool of species across communities. Notably, intermediate levels of internal redistribution sustain high differentiation across a wide range of inflow intensities by allowing local specialization without complete isolation. This perspective emphasizes that spatial differentiation emerges from the balance between diversity input and internal circulation, rather than from dispersal intensity alone.

Our modelling approach allowed us to evaluate how different mechanisms determine the relationship between β-diversity and dispersal, relying only on basic ecological processes (i.e., local recruitment, death, dispersal, colonization, and local selection). This facilitates the interpretation of the model outcomes and contributes to our mechanistic understanding of how dispersal can determine observed, albeit often overlooked, patterns of β-diversity. Furthermore, our approach incorporates factors that are logistically challenging to include in experimental settings. Specifically, we were able to consider a continuous gradient of dispersal intensity rather than a limited number of discrete dispersal levels, and to evaluate the influence of an external regional pool of species on metacommunity dynamics (Grainger and Gilbert 2016). However, some limitations regarding the assumptions of our model should be considered. First, simulations were run on metacommunities containing only a small number of interconnected communities. Nevertheless, different metacommunity structures and directional dispersal influence colonization rates affecting β-diversity differentially among patches (Brown and Swan 2010; Carrara et al. 2012; Altermatt et al. 2013; Seymour and Altermatt 2014; Borthagaray et al. 2025). Second, we did not explicitly include biotic interactions among species which have the potential to strongly affect community structure (Haegeman and Loreau 2014; Soininen et al. 2017; García-Girón et al. 2020; Lu 2021; Ortiz et al, 2023b; Argueta-Guzmán et al. 2025; Souza et al. 2025). Third, we simulated dispersal as a purely stochastic process, although active dispersal may affect biodiversity at several scales (Ye and Wang 2023). Finally, we focused on a specific type of priority effects based on niche preemption in which pioneer species only reduced the amount of resource available (i.e., space) to later dispersers. However, priority effects based on niche modification, where early-arriving species modify local environmental conditions should also be considered (Fukami 2015). The theoretical and empirical analysis of these factors represents promising avenues for future research that may reveal novel mechanisms underlying the assembly process, function and stability of natural systems. We highlight the need for a better understanding of the relative contribution–and interaction–of stochastic versus niche-mediated determinants of β-diversity in metacommunities (Catano et al. 2017; Arim et al. 2023).

