## Supplementary Material for "The interaction of dispersal, species sorting, and priority effects, explains a unimodal response of β-diversity to immigration rates"

**ORCID**

E.O.: 0000-0002-1025-6742

R.R.J.: 0000-0002-0108-7502

M.A.: 0000-0002-7648-8909

### Supplementary Material

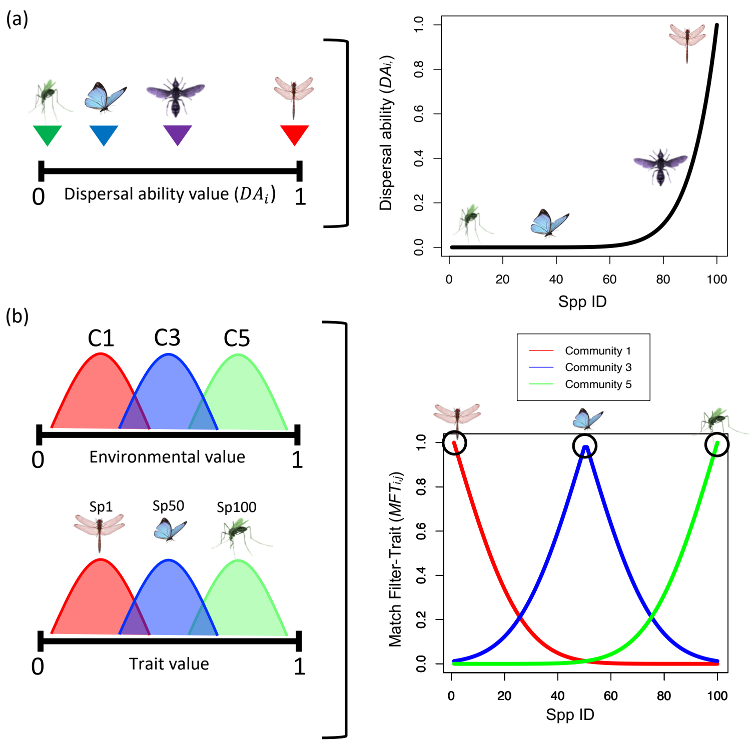


**Figure S1. Determination of species dispersal ability (a) and species local performance (b). (a) Species dispersal ability ($DA_i$) was defined as: $DA_i = x_i^{10*Q}$, where $x_i$ is a value uniformly drawn from the interval [0,1], and $Q$ is a parameter that determines the magnitude of the interspecific differences in $DA_i$. Larger $DA_i$ values determine higher dispersal abilities. (b) Species local performance was determined by assigning to each local community and species an environmental and trait value distribution respectively with mean $\mu$ drawn from the interval [0,1] and constant standard deviation $\sigma$. The local performance of a species $i$ in a local community $p$ was defined as the overlap between the trait distribution of $i$ and the environmental distribution of $j$ (i.e. Match Filter-Trait, $MFT_{i,p}$). If $MFT_{i,p} = 1$, the overlap between the two distributions is maximum which means that $i$ is the species best adapted to community $p$ showing the highest local performance.**

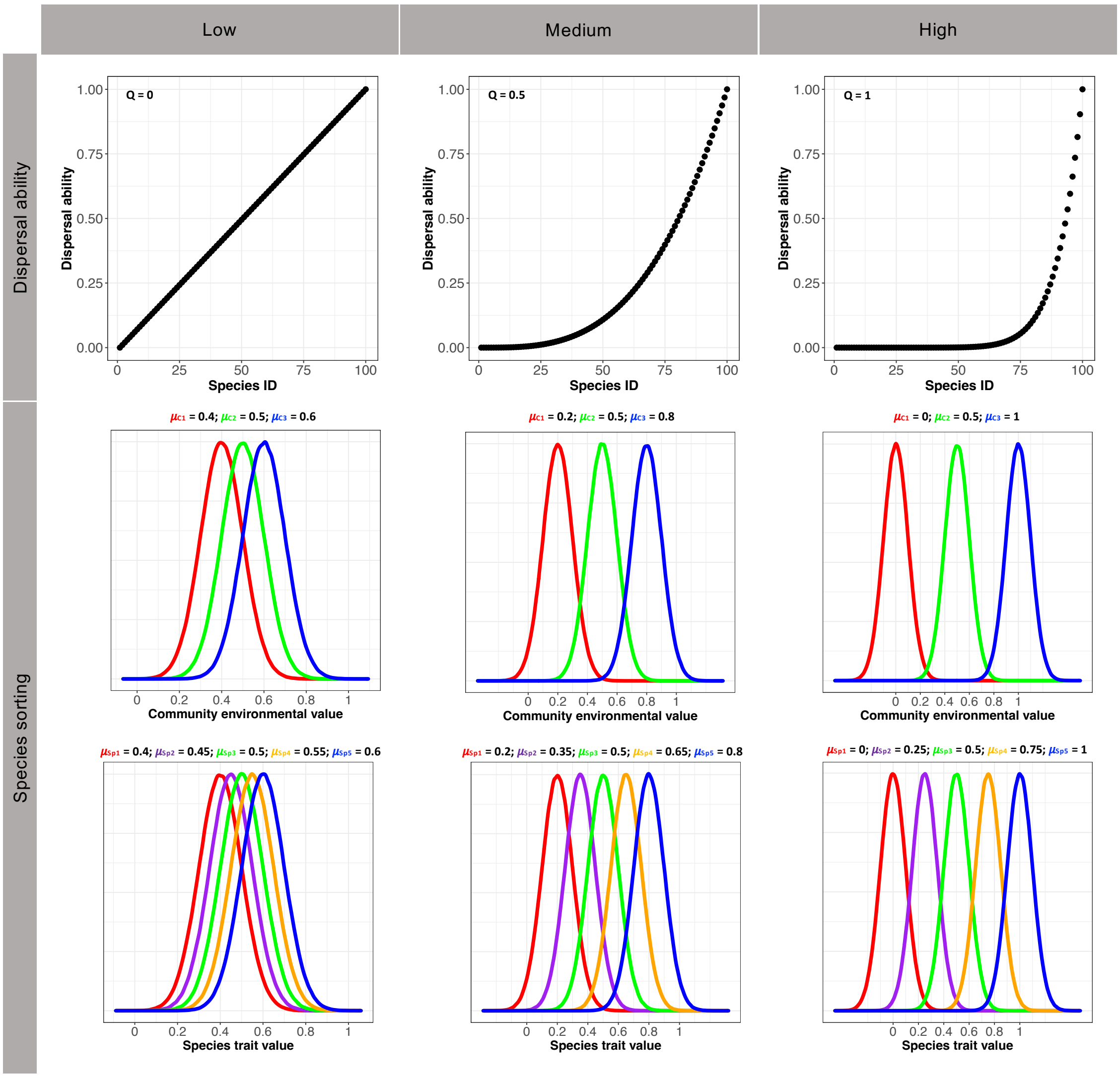


**Figure S2. (a) Variations in interspecific dispersal ability, $DA_i$, were determined by parameter $Q$. Large $Q$ values magnify the differences in dispersal ability between good and bad dispersers. (b) Variations in species local performance were determined by the distance between the mean $\mu$ of each distribution. The longer the distance between the values of $\mu$, the more different local communities and species are in terms of their environmental conditions and traits respectively.**

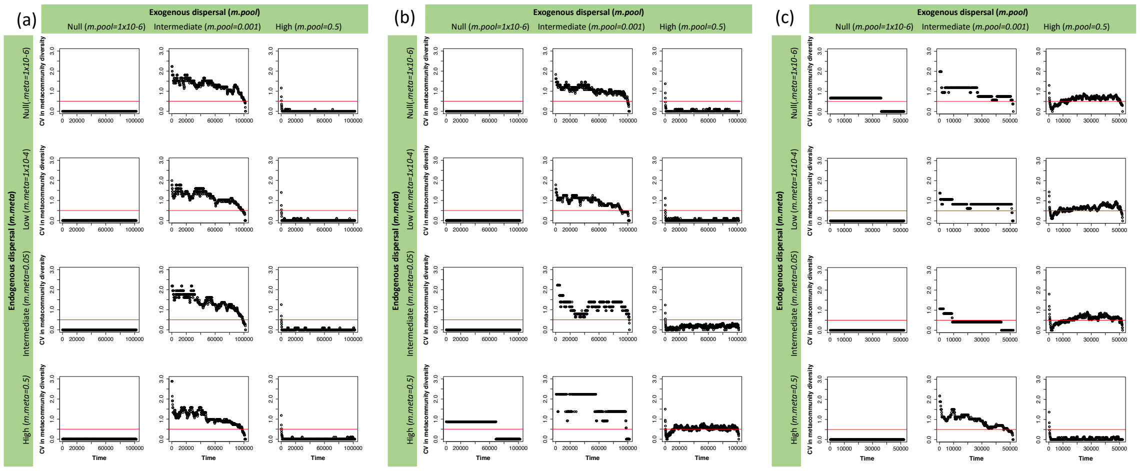


**Figure S3. Variation in metacommunity diversity (coefficient of variation) during the lottery dynamics under the scenarios of (a) random priority effects ($H_1$), (b) priority effects conditioned by species dispersal ability ($H_2$), and (c) species sorting ($H_3$) for different exogenous (m.pool) and endogenous (m.meta) dispersal values. Metacommunity dynamics are considered to be stable when CV values are smaller than 0.5 (red lines).**

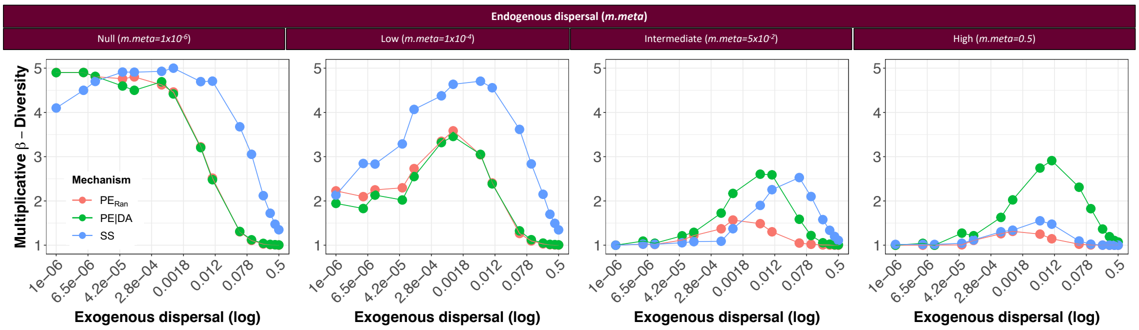


**Figure S4. Trends in β-diversity (multiplicative) along the exogenous dispersal gradient (*m.pool*), considering different intensities of endogenous dispersal (*m.me*ta).**

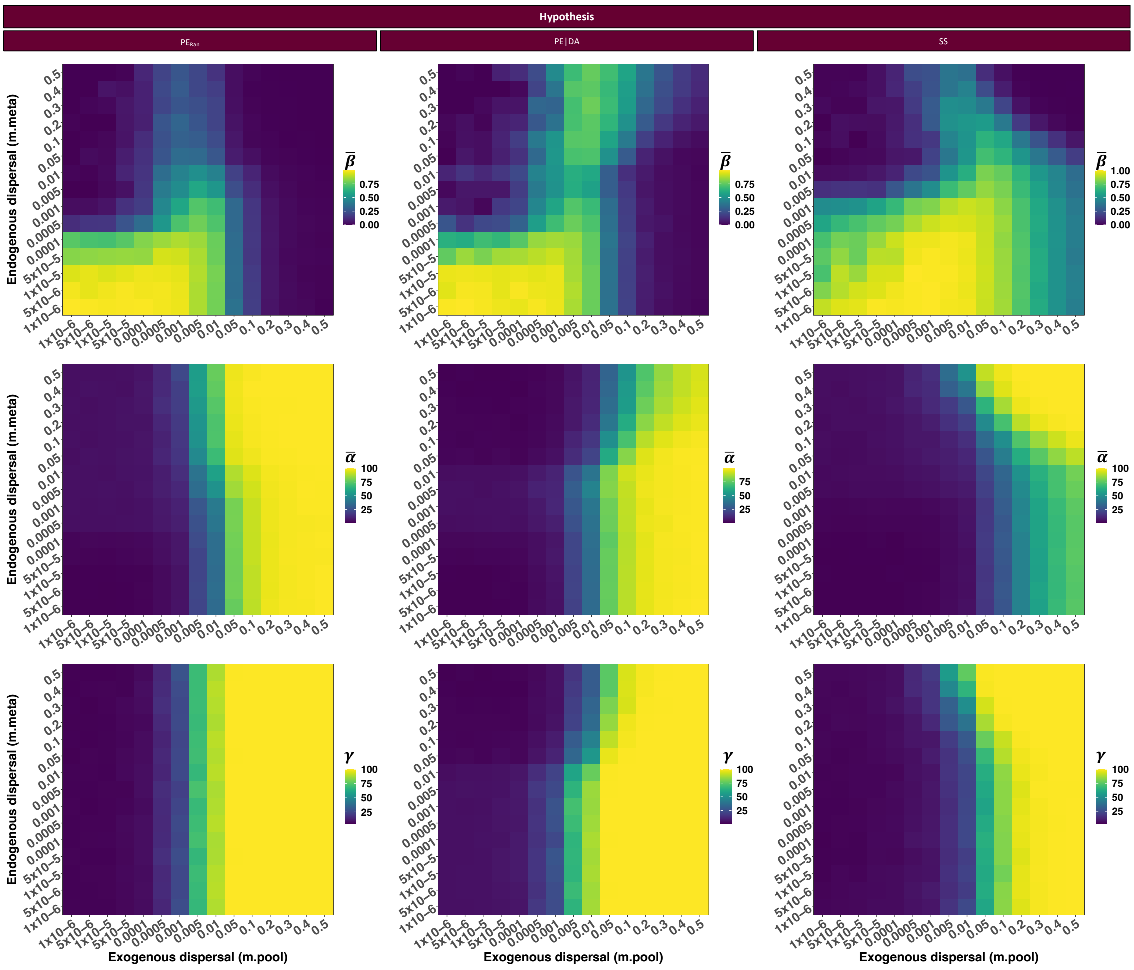


**Figure S5. Mean β-(Jaccard), mean α- and γ-diversity for different combinations of exogenous (m.pool) and endogenous dispersal (m.meta) probabilities.**

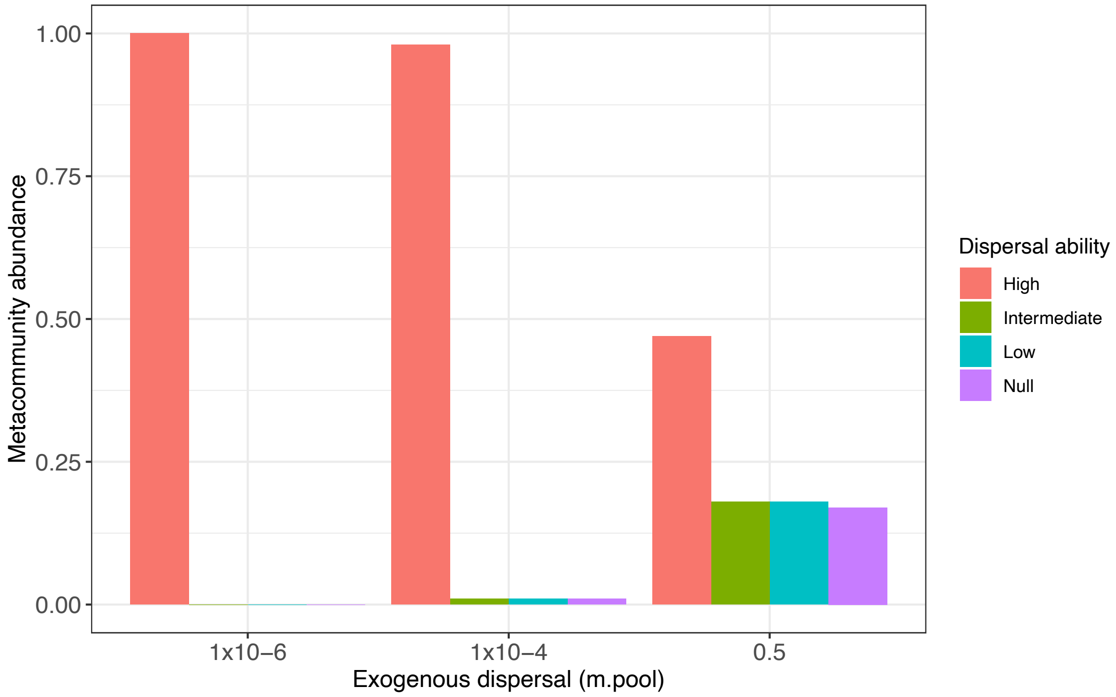


**Figure S6 Metacommunity abundance of species with contrasting dispersal abilities for three exogenous dispersal probabilities (m.meta=5x10$^{-2}$).**

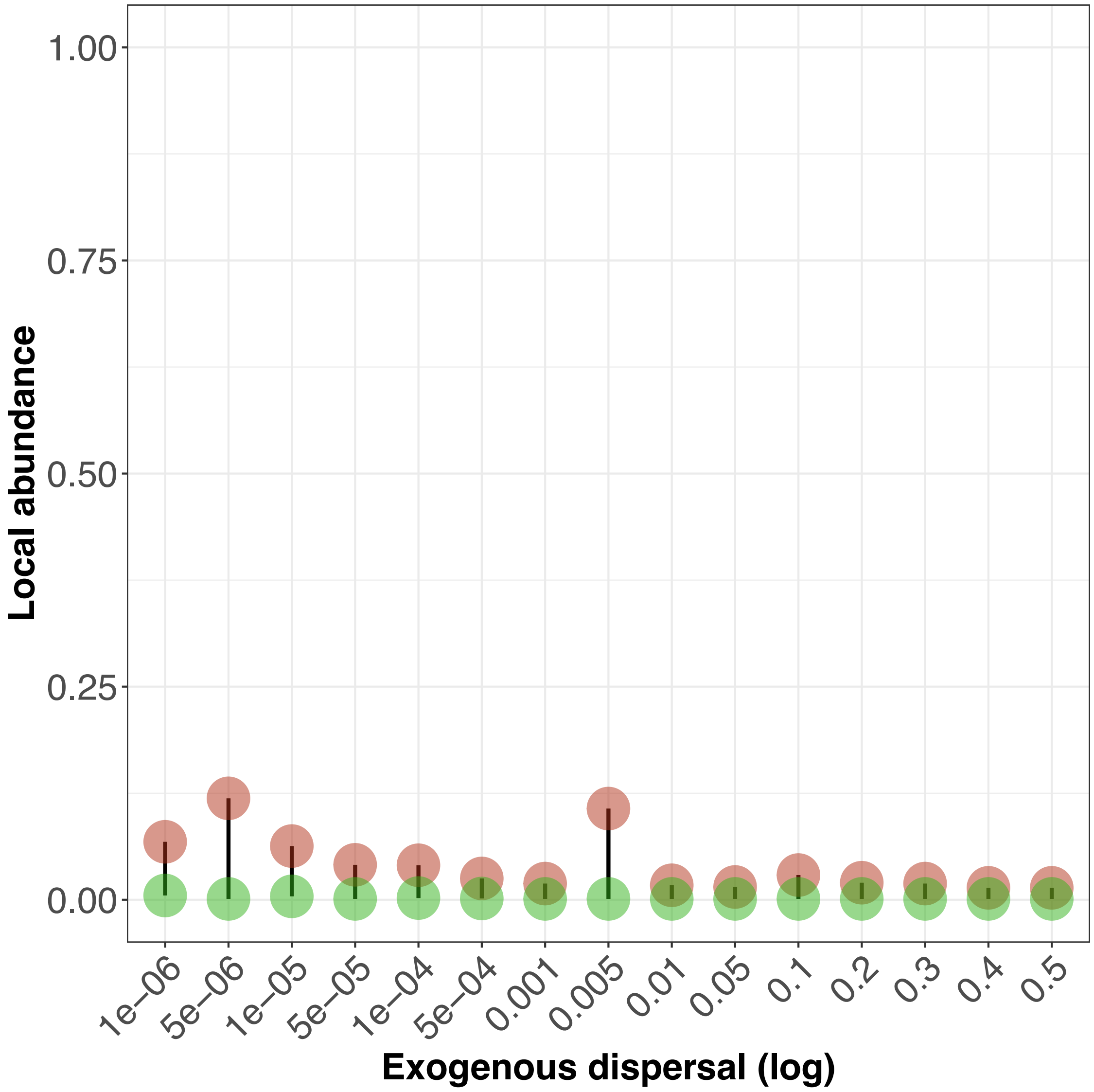


**Figure S7 Effect of species arrival time ($AT_{i,p}$) on local asymptotic species abundance across a gradient of exogenous dispersal rates (*m.pool*), at high endogenous dispersal rates (*m.meta* = $5 \times 10^{-2}$), under the species sorting scenario ($H_3$). Red and green symbols represent species in the earliest and latest 1% of arrivals.**
